# Bimodal 1/*f* noise and anticorrelation between DNA-Water and DNA-Ion energy fluctuations

**DOI:** 10.1101/2022.12.13.520228

**Authors:** Saumyak Mukherjee, Sayantan Mondal, Biman Bagchi

**Affiliations:** Theoretical Chemistry, Ruhr University Bochum, 44801 Bochum, Germany; Department of Chemistry, Boston University, Boston, Massachusetts 02215, United States; Solid State and Structural Chemistry Unit, Indian Institute of Science, Bangalore 560012, India

**Keywords:** DNA, Energy fluctuations, 1/*f* noise, coupled fluctuations, dielectric relaxation, solvation dynamics

## Abstract

Coupling between the fluctuations of DNA and its surroundings consisting of water and ions in solution remains poorly understood and relatively less investigated as compared to proteins. Here, with the help of molecular dynamics simulations and statistical mechanical analyses, we explore the dynamical coupling between DNA, water, and counterions through correlations among respective energy fluctuations in both double (ds-) and single-stranded (ss-) DNA solutions. Fluctuations in the collective DNA-Water and DNA-Ion interaction energies are found to be strongly anti-correlated across all the systems. The fluctuations of DNA self-energy, however, are weakly coupled to DNA-water and DNA-ion interactions in ds-DNA. An enhancement of the DNA-water coupling is observed in ss-DNA, where the system is less rigid. All the interaction energies exhibit 1/*f* noise in their energy power spectra with surprisingly prominent bimodality in the DNA-water and DNA-ion fluctuations. The nature of the energy spectra appears to be indifferent to the relative rigidity of the DNA. We discuss the role of the observed correlations in ion-water motions on a DNA duplex in the experimentally observed anomalous slow dielectric relaxation and solvation dynamics, and in furthering our understanding of the DNA energy landscape.

**TOC GRAPHIC:** 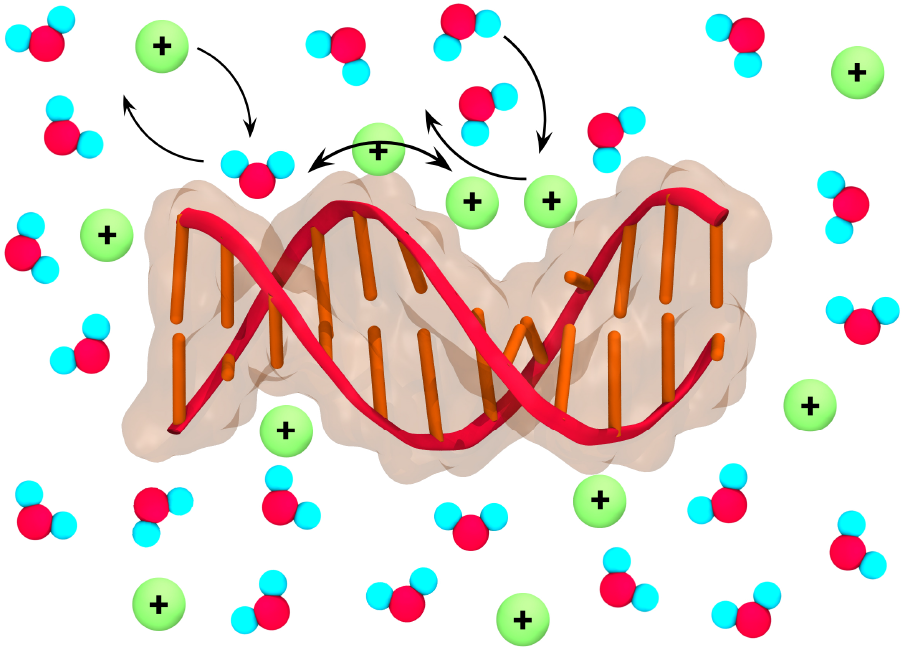

## I. INTRODUCTION

The functional role of water in biology continues to be intensely studied and debated. Although water is recognized as the matrix or elixir of life, the precise microscopic mechanism of its role remains elusive. Because of the small size and low molecular weight, water molecules are continuously in a state of rapid motion, even within biological cells.^1,2^ The structure and dynamics of these biological waters are substantially different from neat bulk water.^3,4^ It has been widely recognized that water molecules present in biological systems act as active components in a multitude of processes such as bond-breaking events, electron transfer reactions, and also in conformational changes.^2–6^ Moreover, water has been shown to modulate internal fluctuations within biomolecules.^7,8^ Although the precise origin of this controlling effect of water remains largely unknown, the conformational and energy fluctuations in water are believed to be major drivers that help modulate the coupling with proteins.^9^

Recent studies have explored some aspects of the functional role of water in proteins following the intriguing observations of Frauenfelder that suggest “water slaves proteins”.^9,10^ Such investigations have revealed strong signatures of coupling between water and protein fluctuations both in collective energies and in forces acting on specific bonds in the proteins. This aspect of coupled fluctuations, however, remains unmapped in the case of other biomolecules, such as DNA.

Energy fluctuations, particularly correlations among different contributing energy components in the system, can unveil crucial insight into the activity of a biomolecule. Analogous to the bond-breaking events in proteins, in a double-stranded (ds-) DNA, one would like to know how these fluctuations affect elementary processes such as the dissociation of the hydrogen bonds between the base pairs. Many important biological processes involve the separation of the ds-DNA locally, and also bubble formation.^11,12^ These are slow processes and involve substantial energy costs. The mechanism of the flow of energy to and from the reaction centers, however, is largely unknown. Most importantly, the role of water and counterions, which are major crowders in the vicinity of the DNA are ill-understood. Large fluctuations in the position and orientation of surrounding water molecules and counterions could be responsible for the initiation of processes such as the separation of the helices in a ds-DNA, for example. In the case of drug-DNA intercalations, studies have shown that the drug initially binds to the minor groove. Any subsequent rearrangement that would involve crossing an energy barrier must again be driven by energy fluctuations of ions and water.^13^

A study of correlations among energy fluctuations can provide information about the DNA energy landscape as well as the origin of the experimentally observed slow solvation dynamics and dielectric relaxation. It is noteworthy that these fluctuations contain both slow, intermediate, and fast components that are not easy to perceive. Moreover, the presence of counterions adds complexity to the energy landscape. There exist several studies that shed light on the dynamics of ds-DNA in aqueous solutions. Extensive experimental studies, aided by computer simulations, reveal that the solvation dynamics of ds-DNA show vastly different properties as compared to proteins.^14–17^ Most interestingly, energy fluctuations in DNA exhibit an ultra-slow long-time decay (of the order of hundreds of nanoseconds).^15^ The long-time part of the decay can be characterized by a power law, with a small fractional exponent. Such dynamical characteristics, as seen in DNA, are absent in proteins and suggest that the nature and the extent of coupling of a DNA duplex with its environment (water and counterions) are different from that of a protein. At the same time, it is also pertinent to interrogate the same energy correlations for the single-strand DNA (ss-DNA). Here the biopolymer is more exposed to its aqueous environment and capable of executing large-amplitude conformational fluctuations on account of reduced structural rigidity.

An earlier group of studies, pioneered by Oosawa, investigated the dielectric relaxation (DR) of aqueous DNA solutions.^17^ These studies revealed that the dielectric relaxation was markedly non-Debye, especially in the low-frequency side of the Cole-Cole plot. It was proposed that the motion of counterions along the rigid ds-DNA backbone could be responsible for such anomalous dielectric relaxation. While the origin of this non-Debye relaxation is still not clear, it seems conceivable that the slow power law solvation dynamics and the anomalous dielectric relaxation are related. This was proposed by the continuum model theories of solvation that use the dielectric function as a measure of polarization relaxation.^18^ The obvious questions that follow are:

- What are the reasons for such slow, anomalous dynamics in dielectric relaxation and solvation time correlation function?
- Are the motions of water and ions correlated? A recent study employed the method of continuous time random walk to study the motions of ions and water on the DNA backbone.^16^ However, a detailed connection of the counterion motions to DR was not attempted.
- Are the internal motions of DNA at all affected by the fluctuations of water, like in proteins?
- Moreover, do the counterions that diffuse along the phosphate backbone of a DNA have any influence?
- How do all these motions affect the functions of DNA?

To investigate these issues, we simulate two 12 base-pair (BP) long DNAs, one with an abundance of Guanine-Cytosine (GC) BPs and the other being rich in Adenine-Thymine (AT) BPs. To probe dynamic correlations between DNA, water and ions we monitor the correlations among fluctuation in the interaction energies among these components of the solution.

It has been recently shown that the fluctuations of self-energy in proteins and those in protein-water interactions are strongly anti-correlated.^7^ This points towards a flow of energy between protein and water that provides a channel for the solvent control of protein fluctuations, a hypothesis, as already mentioned, put forward by Frauenfelder and co-workers.^9^ However, the structural differences between proteins and DNAs make it unclear whether such an effect is also present in the latter case. Unlike protein, DNA has a regular arrangement of the BPs, with strong hydrogen bonds (HBs) between them. While the AT pair has two HBs, the GC pair has 3 HBs. Hence, DNA does not have the structural flexibility that is enjoyed by a protein. The rigidity of the DNA structure is further reinforced by the *π*-stacking interactions between the aromatic rings in the neighboring bases, and also the repulsive forces acting between the negative charges on the phosphate backbone. The present study also attempts to decipher the consequences of this architectural robustness on the coupling of structural fluctuations in DNA with those in water and counterions. Furthermore, we explore the origins of the experimentally observed anomalous dielectric relaxation and solvation dynamics of DNA in terms of coupled water-ion dynamics on the frame of the DNA phosphate backbone.

## II. COMPUTATIONAL METHODS

All molecular dynamics simulations were performed using the GROMACS 2021.3 simulation package.^19^ The initial structures of the double-stranded AT-rich (*ATATCTAGATAT*)_2_ and GC-rich (*GCGCTCGAGCGC*)_2_ DNAs were prepared using the Sequence to Structure web server (link). These are shown in Fig. 1. For the single-stranded DNA, one of the chains in each of the aforementioned sequences was taken. Each DNA was solvated in *∼*5500 explicit water molecules in a box of dimensions (5*×* 5*×* 7) *nm*^3^. Sodium ions were introduced to neutralize the charges on the DNA. The AMBER99SB-ILDN force field^20^ was used for the simulation of the DNA and Ions, whereas the water molecules were described by the TIP3P model^21^, which is often used along with the aforementioned nucleic acid force field.

**FIG. 1.**
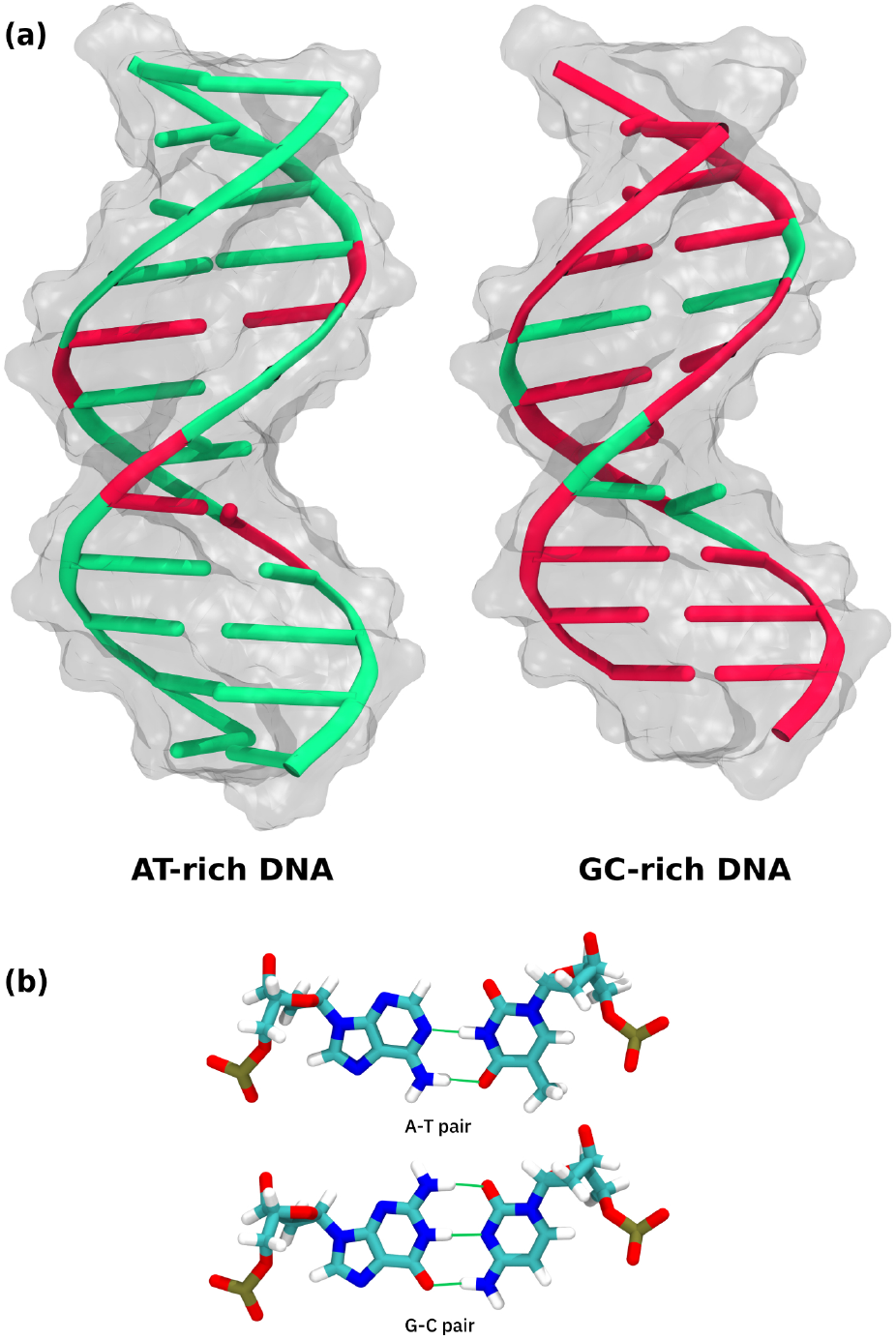
(a) Snapshots of twelve base pair AT-rich and GC-rich double-stranded DNAs studied in this work. The green color represents the A-T pairs, whereas the red color represents the G-C pairs. (b) Hydrogen bonding (HB) between base pairs: two HBs between A and T; three HBs between G and C.

The energy of each system was first minimized using the steepest decent algorithm. The energy-minimized systems were then equilibrated for 500 ps (picosecond) under NpT conditions (pressure (p) = 1 bar and temperature (T) = 300 K) with harmonic position restraints on the DNA atoms. Thereafter, the restraints were removed and further NpT equilibrations of 1 ns (nanosecond) were performed. This was followed by 1 ns NVT equilibrations (T = 300K). The final production runs were carried out for 1 *μ*s (microsecond) under similar NVT conditions. The data were saved at 10 ps intervals.

The Parrinello-Rahman barostat^22^ and Nosé-Hoover thermostat^23,24^ were used to regulate pressure and temperature, respectively, whenever necessary. The Leap-Frog algorithm was used to integrate the equations of motion with a time step of 2 fs (femtosecond). A cut-off radius of 1 nm was used for neighbor searching and for the short-range part of nonbonded interactions. The long-range interactions were taken care of by the Particle Mesh Ewald (PME) method^25^ with a Fourier grid spacing of 1.2 nm. All bonds in the DNAs were constrained using the LINCS algorithms.^26^

To minimize the errors in energy calculations due to the short-range cut-off, we post-process the simulation trajectories using the *mdrun rerun* routine in GROMACS with an increased cut-off value of 2.4 nm for both Coulomb and van der Waals interactions. The individual energy contributions from DNA, water, and ions, and their cross-interactions are computed by using the *energygrps* option in GROMACS.

## III. RESULTS AND DISCUSSION

### A. Energy Fluctuations in ds-DNA

The total energy fluctuations of a DNA solution (*δ E*) can be decomposed based on the components in the system, namely, the DNA, water, and counterions, along with their cross-interaction terms [Eq. 1].

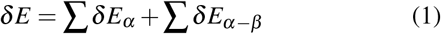

The first term in Eq. 1 denotes the self-interactions (*δ E*_*D*_, *δ E*_*W*_, and *δ E*_*I*_; where *D, W*, and *I* represent DNA, water and ions respectively) and the second one denotes the cross-terms (*δ E*_*D−W*_, *δ E*_*D−I*_, and *δ E*_*W*−*I*_).

In Fig. 2, we plot the time evolution of pairs of energy fluctuations in the AT-rich ds-DNA system. As seen in Fig. 2a, *δ E*_*D*_ and *δ E*_*D−W*_ are apparently uncorrelated. Similar behavior is seen for *δ E*_*D*_ and *δ E*_*D*_−_*I*_ in Fig. 2b. However, a strong anti-correlation can be observed between *δ E*_*D−W*_ and *δ E*_*D−I*_ in Fig. 2c. GC-rich ds-DNA shows similar energy fluctuation behavior. This observation begs quantification of the extent of correlation between these pairs of energy fluctuations.

**FIG. 2.**
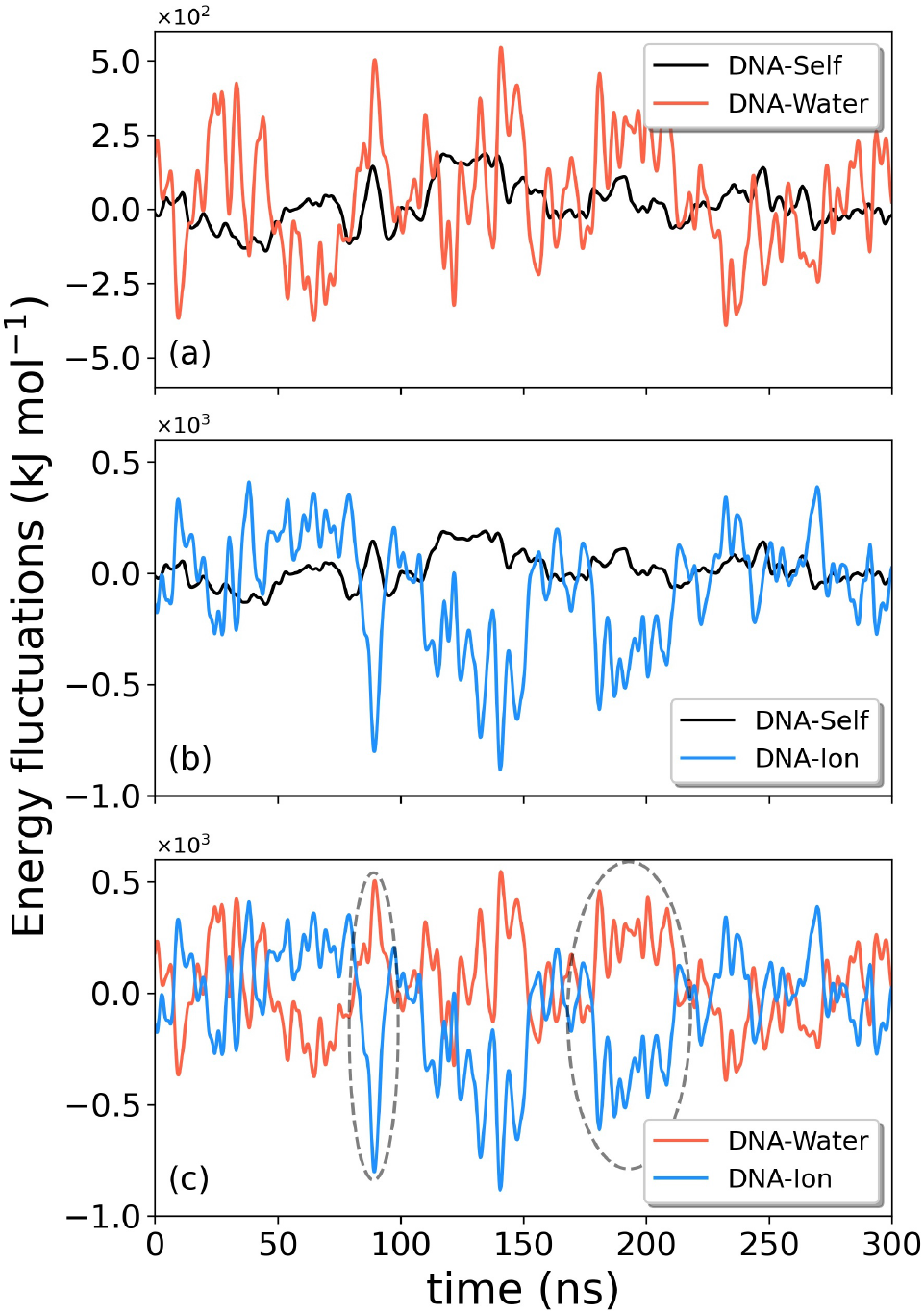
Time traces of energy fluctuations in AT-rich ds-DNA system shown for a period of 300 ns. The colors black, red, and blue respectively represent DNA-self (*δ E*_*D*_), DNA-Water (*δ E*_*D−W*_), and DNA-Ion (*δ E*_*D−I*_) energy fluctuations. The plots are smoothed using Gaussian averaging for clarity of representation. The grey ellipses are examples of anti-correlated fluctuation events between *δ E*_*D−W*_ and *δ E*_*D−I*_.

To quantify the extent of coupling among these energy fluctuations terms, we compute the respective Pearson correlation coefficients (*ρ*_*xy*_ = *cov*(*x, y*)*/*(*s*_*x*_*s*_*y*_), where *cov*(*x, y*) is the co-variance between the two variables x and y, and *s*_*x*_ and *s*_*y*_ are their standard deviations) for both the AT-rich and the GC-rich DNA systems. The obtained values are presented in Table I.

**TABLE I.** Pearson correlation coefficients between the energy fluctuation terms in aqueous ds-DNA. The highly anticorrelated terms are shown in boldface.

| Energy term<br>($x$) | Energy term<br>($y$) | $\rho_{xy}$<br>(AT-rich) | $\rho_{xy}$<br>(GC-rich) |
| --- | --- | --- | --- |
| $\delta E_D$ | $\delta E_{D-W}$ | -0.21 | -0.16 |
| $\delta E_D$ | $\delta E_{D-I}$ | -0.15 | -0.22 |
| $\delta E_{D-W}$ | $\delta E_{D-I}$ | **-0.85** | **-0.83** |

*δ E*_*D*_ shows 21 % and 16 % anticorrelation with *δ E*_*D − W*_ in the AT-rich and the GC-rich systems respectively. These values are substantially lower than the analogous correlation coefficients observed in protein systems (*∼*90 %).^5^ Similarly, a relatively small anticorrelation is observed between *δ E*_*D*_ and *δ E*_*D*−*I*_. In contrast to this, a substantially strong anticorrelation exists between *δ E*_*D−W*_ and *δ E*_*D*−*I*_ in both the ds-DNA systems. The other possible combinations of energy pairs are irrelevant to the present study and hence omitted in this analysis.

The stark difference between the extent of protein-water and DNA-water anticorrelations shows that, unlike proteins, the dynamics of ds-DNA is relatively decoupled from that of water. This might be a consequence of the increased rigidity of the ds-DNA by virtue of its multiple regularly spaced hydrogen bonds between the base pairs.

Because of the multiple atoms and repetitive structural features involved in the interactions when one considers the whole ds-DNA, there is a significant probability of cancellations of multiple energy fluctuation components that might lead to the observed low anticorrelation values. In order to check this we perform a similar analysis at a local scale for specific base pairs, namely (i) *G*_19_ and *C*_6_ in the GC-rich DNA and (ii) *A*_17_ and *T*_6_ in the AT-rich DNA. The Pearson correlation coefficients are provided in Table II. We find that the trend that was observed on a global length scale is preserved in local energy fluctuations as well. The anti-correlation between base pair-water and base pair-ion is approximately 0.8– 0.9. Other pair interaction energies are found to be weakly correlated.

**TABLE II.** Pearson correlation coefficients between energy fluctuation terms of specific tagged base pairs in (a) GC-rich and (b) AT-rich ds-DNA solution. The highly anticorrelated terms are shown in bold-face.

| AT-rich |  |  | GC-rich |  |  |
| --- | --- | --- | --- | --- | --- |
| Energy (x) | Energy (y) | $\rho_{xy}$ | Energy (x) | Energy (y) | $\rho_{xy}$ |
| $A_{17}$-DNA | $A_{17}$-Water | -0.09 | $G_{19}$-DNA | $G_{19}$-Water | -0.01 |
| $A_{17}$-DNA | $A_{17}$-Ion | -0.17 | $G_{19}$-DNA | $G_{19}$-Ion | -0.14 |
| **$A_{17}$-Water** | **$A_{17}$-Ion** | **-0.82** | **$G_{19}$-Water** | **$G_{19}$-Ion** | **-0.89** |
| $T_6$-DNA | $T_6$-Water | -0.06 | $C_6$-DNA | $C_6$-Water | 0.00 |
| $T_6$-DNA | $T_6$-Ion | -0.12 | $C_6$-DNA | $C_6$-Ion | -0.11 |
| **$T_6$-Water** | **$T_6$-Ion** | **-0.88** | **$C_6$-Water** | **$C_6$-Ion** | **-0.87** |

The high degree of anti-correlation between the ions and water denotes that in a configurational space description, the ions and water molecules can exchange positions. Our simulation systems do not have buffer ions. Hence, the cross-anticorrelation comes entirely from the counterions that balance the charges on the DNA phosphate backbone. The dynamics of these counterions along the double-helical backbone play a crucial role in determining the solvation dynamical properties of DNA solutions. It was recently proposed that such motion of ions, which resembles the continuous random walk model proposed by Montroll-Scher-Lax, may be responsible for the power law behavior of DNA solvation dynamics. The rigid structure of the DNA itself provides a platform for the ions to move around.

However, do the ions and water indeed show an exchange dynamics? To answer this, we monitor the local environmental density fluctuations around the phosphate groups in our simulations. Interestingly, the number of ions and that of water molecules within 0.3 nm of the phosphate groups show an anticorrelation with a coefficient of -0.33. This clearly shows that on several occasions, water molecules replace the counterions on the DNA backbone.

To perceive the origin of the correlation coefficients given in Table I, we need to understand the timescales associated with the relaxation of these energy fluctuations. Toward that goal, we compute their autocorrelation functions (ACF) (*C*_*E*_ (*t*)), as shown in Fig. 3. *C*_*E*_ (*t*) is calculated for energy fluctuations in both AT- and GC-rich ds-DNA systems.

**FIG. 3.**
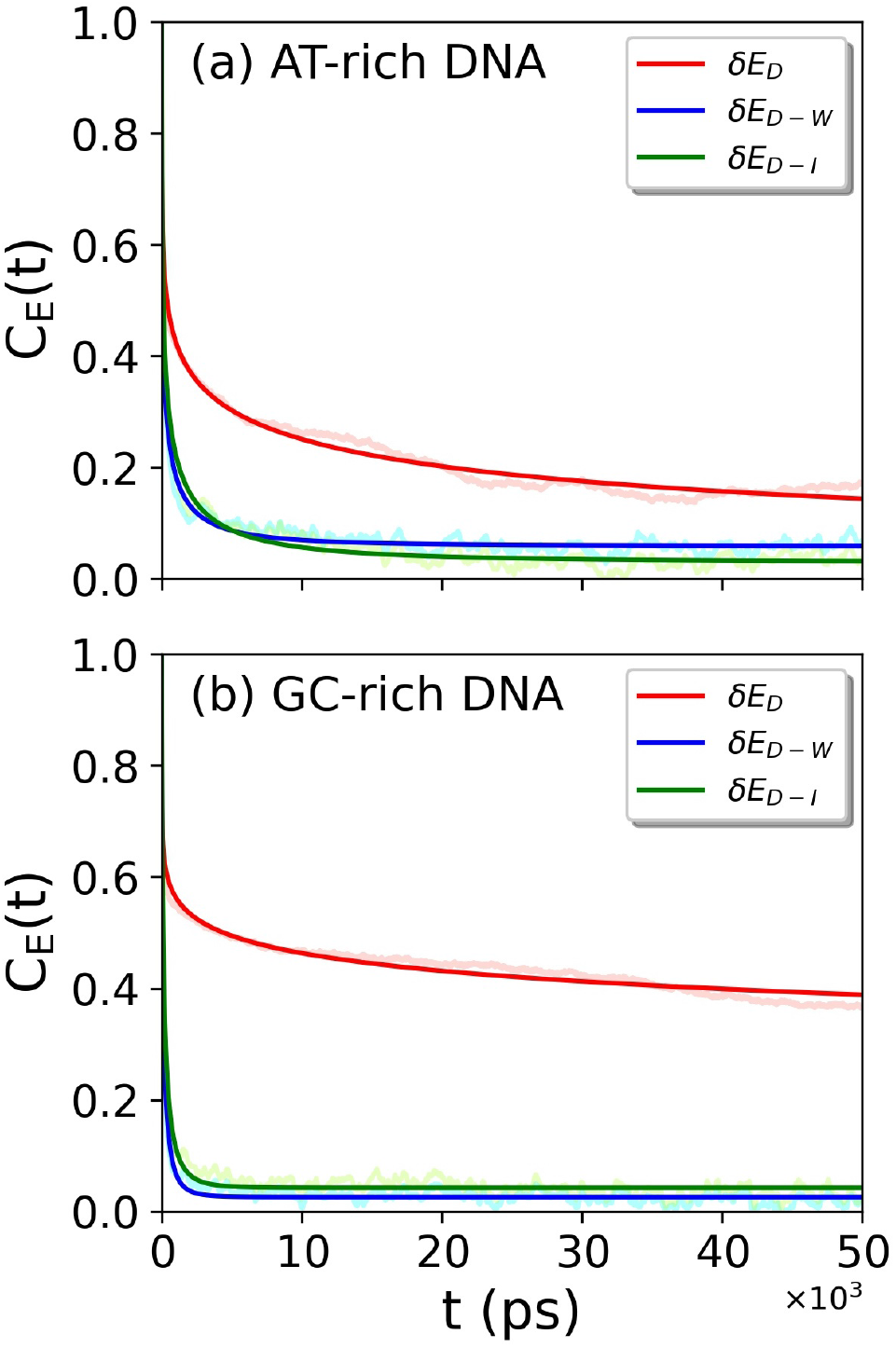
Autocorrelation functions (ACFs) of energy fluctuations in double-stranded DNA. These are shown for both (a) AT- and (b) GC-rich systems. The red, blue, and green colors represent ACFs of DNA-self, DNA-water, and DNA-ion interaction energy fluctuations respectively.

The ACFs in Figure 3 are fitted to triexponential functions 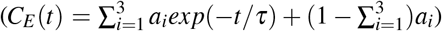 and the data are presented in Table III.

**TABLE III.**
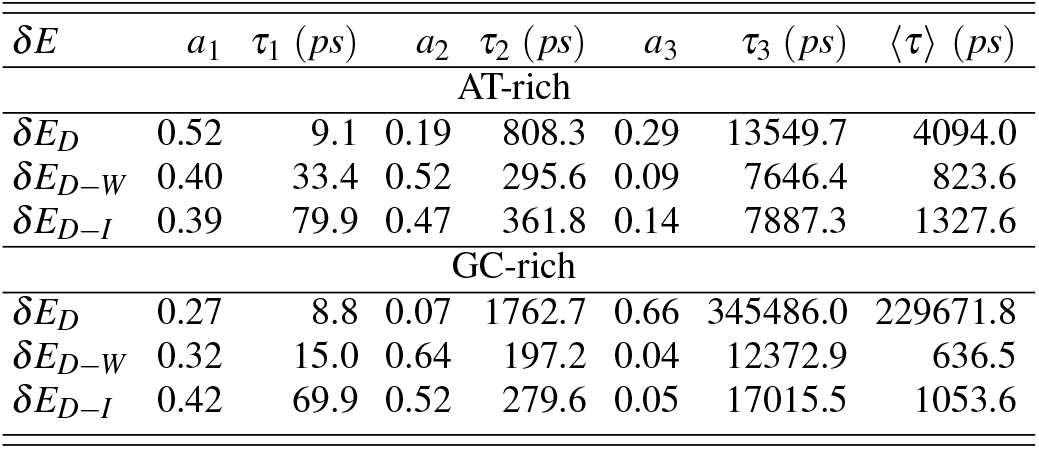
Triexponential fitting parameters of energy ACFs of aqueous ds-DNA system.

| $\delta E$ | $a_1$ | $\tau_1$ (ps) | $a_2$ | $\tau_2$ (ps) | $a_3$ | $\tau_3$ (ps) | $\langle \tau \rangle$ (ps) |
| --- | --- | --- | --- | --- | --- | --- | --- |
| AT-rich |  |  |  |  |  |  |  |
| $\delta E_D$ | 0.52 | 9.1 | 0.19 | 808.3 | 0.29 | 13549.7 | 4094.0 |
| $\delta E_{D-W}$ | 0.40 | 33.4 | 0.52 | 295.6 | 0.09 | 7646.4 | 823.6 |
| $\delta E_{D-I}$ | 0.39 | 79.9 | 0.47 | 361.8 | 0.14 | 7887.3 | 1327.6 |
| GC-rich |  |  |  |  |  |  |  |
| $\delta E_D$ | 0.27 | 8.8 | 0.07 | 1762.7 | 0.66 | 345486.0 | 229671.8 |
| $\delta E_{D-W}$ | 0.32 | 15.0 | 0.64 | 197.2 | 0.04 | 12372.9 | 636.5 |
| $\delta E_{D-I}$ | 0.42 | 69.9 | 0.52 | 279.6 | 0.05 | 17015.5 | 1053.6 |

Table III suggests that the energy autocorrelations are accompanied by three timescales, namely, (i) a fast sub 100 ps timescale, (ii) an intermediate timescale, and (iii) a substantially slow component. The decay of the time correlation function for DNA-self energy fluctuation in GC-rich DNA is slower than that of AT-rich DNA. This can be attributed to the enhanced rigidity of the GC-rich DNA due to more HBs between complementary base pairs. The enhanced rigidity makes the dynamical fluctuations slower. On the other hand, the DNA-Water and DNA-Ion autocorrelations decay relatively faster for GC-rich DNA, compared to the same in AT-rich DNA. This can again be understood by the enhanced rigidity of the GC-rich DNA that suppresses the extent of coupling between DNA and water/ion fluctuations.

Fourier transform of the energy autocorrelation functions yields the energy spectra. If the spectrum in log-log representation has a linear slope between 0.5 and 1.5, it is referred to as 1/*f* noise, where *f* stands for frequency. This noise is omnipresent in nature. It has been previously shown that the energy spectra of both water and proteins exhibit 1/*f* noise characteristics with slopes of *∼*0.6 and *∼*0.9 respectively.^27,28^ Recent studies have shown that the total energy (self-energy plus interaction energy with water) spectrum of solvated proteins has bimodal 1/*f* noise features, that originate from the influence of water on protein dynamics.^7^ The two slopes observed are close to what was previously observed for protein and water. In a similar fashion, to understand the influence of water fluctuations in the energy spectrum of DNA, we plot the same in Fig. 4.

**FIG. 4.**
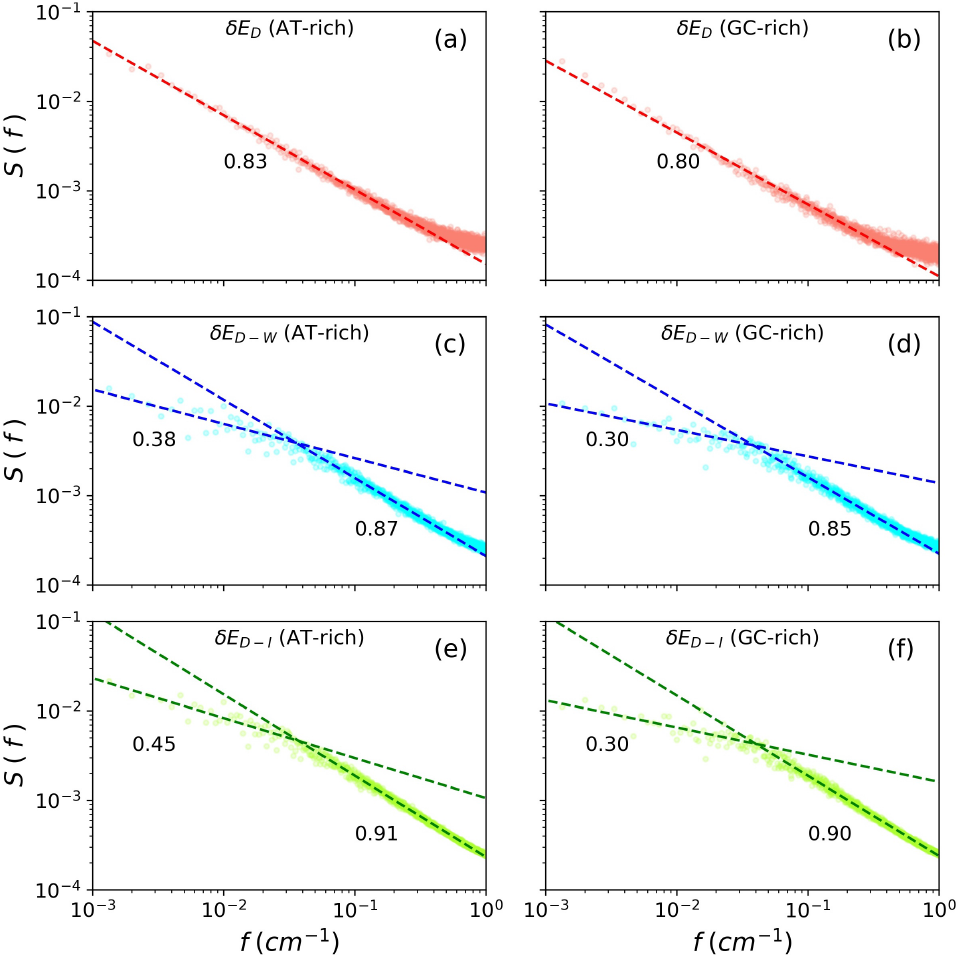
Spectra of the energy fluctuations in ds-DNA solution (log-log representation). (a) DNA-self energy for AT-rich, (b) DNA-self energy for GC-rich, (c) DNA-Water cross interaction energy for AT-rich, (d) DNA-Water cross interaction energy for GC-rich, (e) DNA-Ion cross interaction energy for AT-rich, and (f) DNA-Ion cross interaction energy for GC-rich DNA systems. The numbers denote the slopes of the adjacent lines.

Fig. 4 shows that all the energy terms have 1/f noise-characterized spectra. While the pure self-terms are unimodal, the cross-interaction terms are bimodal, which is similar to what was recently reported for proteins. In proteins, the bimodal protein-water energy spectrum is associated with a high anti-correlation in this coupling interaction. In DNA however, the bimodality exists in spite of a dramatically reduced anti-correlation, as discussed before. This hints that the nature of the energy spectrum is, to a large extent, not influenced by the dynamic behavior of the energy fluctuations. Furthermore, the rigidity of the DNA structure does not have any apparent effect on the slope of the energy spectra. The two gradients in the coupled terms seem to originate from DNA and water/ion. There are several interesting points in this context that deserve special mention and further analysis. It is interesting to note that exponents in the 1/*f* noise spectrum that describe water and ion contributions to DNA energy fluctuation have nearly the same values. This raises an interesting possibility that ions are slaved to water dynamics. In order to resolve the origin, we need to study a pure water-ion system, like an electrolyte solution, which is out of the scope of the present work, but definitely warrants deeper analysis in the future. We further note that the exponent of the water contribution has a value that is substantially lower than that in pure bulk water. This was not the case for protein solutions where the exponents remained almost invariant, thus making it easy to identify the source of energy fluctuations. The smaller value of the exponent for water, signifying a higher degree of correlation, could derive contributions from the two grooves where water molecules are more constrained.

Furthermore, the value of the exponent (*∼*0.85) for DNA self-energy is surprisingly similar to the value (*∼* 0.95) we observed for protein self-energy. The origin of this nearly perfect 1/*f* noise is not clear yet. Ohmine *et al*. originally attributed the 1/*f* noise in the bulk water energy spectrum to intermittency. Subsequently, however, the alternative interpretation in terms of highly collective slow dynamics that gives rise to a stretched exponential-type decay was favored. Mathematically, both can provide 1/*f* noise spectrum, although the mechanisms are rather different. This calls for a study of higher-order correlation functions.

The question that naturally arises is the relationship of the 1/*f* noise to dielectric relaxation and solvation dynamics. Both phenomena display well-known anomalous slow dynamics that are characterized by fractional time or frequency exponents. However, no satisfactory explanation has yet emerged. From the analysis thus presented, one realizes that the structural robustness of DNA, although does not perturb the slopes of the energy spectrum, decouples the DNA and water/ion fluctuations to a great extent. This, in turn, allows a strong coupling between the water and the ions, which results in strong anticorrelated fluctuations. To investigate this aspect in detail, we perform similar analyses on single-stranded (ss-) DNAs (both AT- and GC-rich). ss-DNAs are devoid of the rich HB pattern that is enjoyed by ds-DNA. Hence, it is expected that the structure becomes relatively flexible, which allows us to contest our inferences obtained from the investigations on the more rigid ds-DNAs.

### B. Energy Fluctuations in ss-DNA

As discussed in the previous section, the structural rigidity of the double-helical DNA could be the primary reason that we do not observe a strong anti-correlation between DNA-self and DNA-water cross interactions, as we do in the case of proteins. It is, however, noteworthy that a weak, non-negligible anti-correlation is still although substantially lower than that observed in the case of protein solutions.

To compare the flexibility of the ds- and ss-DNAs, we calculate their root mean square displacements (RMSD). In Fig. 5, we plot the RMSD of atomic positions in ds- and ss-DNA for the GC-rich case. Clearly, the mean RMSD of the latter is substantially greater, which denotes increased structural flexibility. Similar behavior is observed in the case of the AT-rich DNA chains as well.

**FIG. 5.**
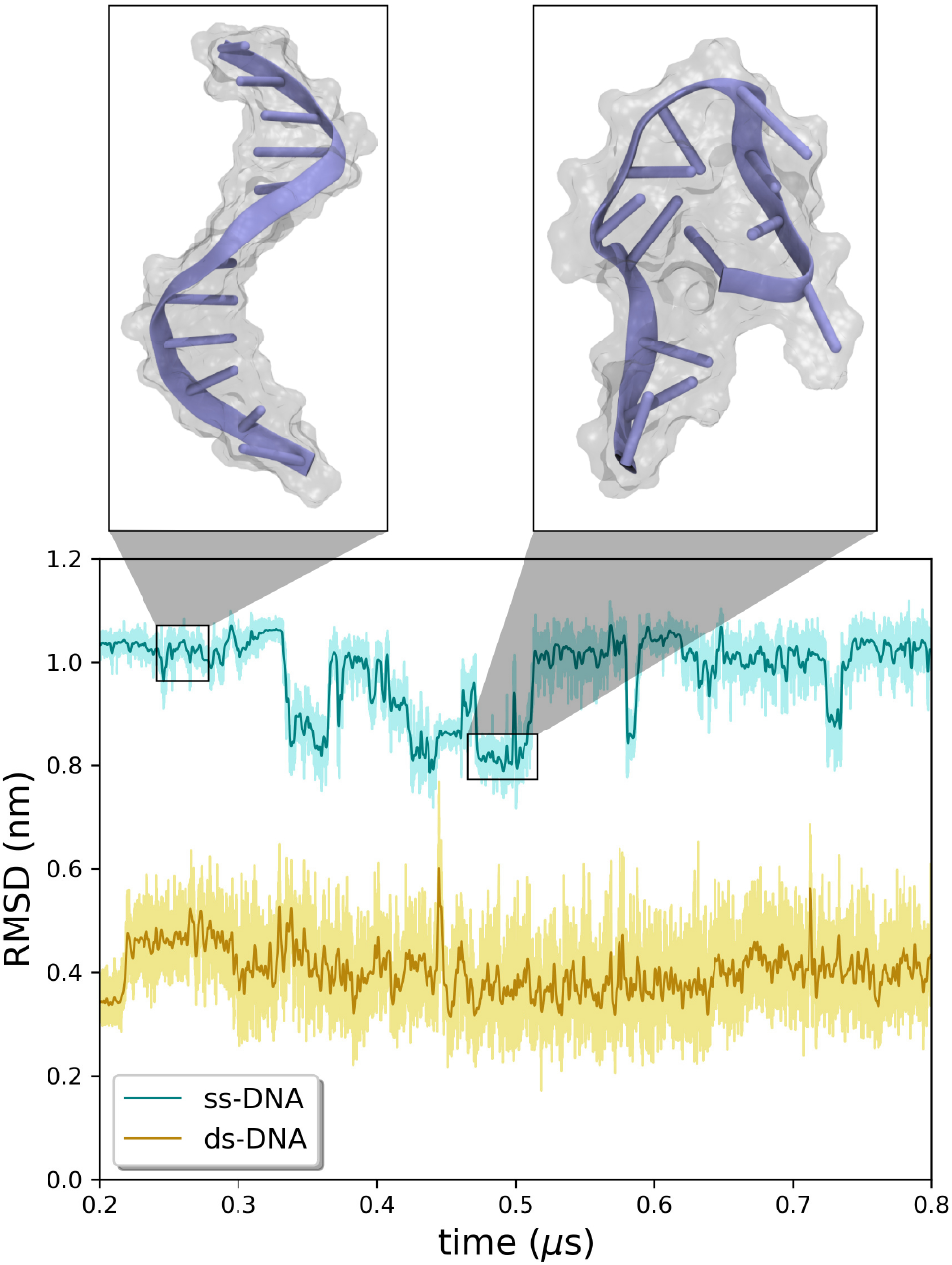
Root mean square displacement (RMSD) of atomic positions in ss- and ds-DNA in the GC-rich system. The darker lines denote the running averages over 1000 data point windows. The RMSD of ss-DNA is greater than that of ds-DNA, which denotes greater conformational fluctuations in the former resulting in reduced rigidity of the structure. These fluctuations may lead to large-scale changes in the structure of the ss-DNA, as shown at two points in the trajectory with representative snapshots of the ss-DNA.

Interestingly, the time traces of RMSD for ss-DNA exhibits several instances of sudden jumps, unlike the ds-DNA. Further probe into the structural states of the ss-DNA before and after such jumps clearly shows substantial structural changes (Fig. 5). This demonstrates that in the absence of base-pair hydrogen bonds (as in ds-DNA) the ss-DNA has increased degrees of freedom resulting in a less rigid phosphate backbone. Consequently, this can be efficiently employed to compare the results obtained in case of the ds-DNA systems.

Table IV provides the Pearson correlation coefficients in the ss-DNA systems for the same energy fluctuation terms as presented in Table. We observe a dramatic increase in the anticorrelation between *δ E*_*D*_ and *δ E*_*D−W*_. On the other hand, *δ E*_*D*_ and *δ E*_*D−I*_ become even less anti-correlated. These two observations speak in favor of the strong anticorrelation between *δ E*_*D*−*W*_ and *δ E*_*D*−*I*_, which is retained in the ss-DNA systems as well.

**TABLE IV.** Pearson correlation coefficients between energy fluctuation terms in aqueous ss-DNA. The highly anticorrelated terms are shown in boldface.

| Energy term<br>(x) | Energy term<br>(y) | $\rho_{xy}$<br>(AT-rich) | $\rho_{xy}$<br>(GC-rich) |
| --- | --- | --- | --- |
| $\delta E_{D-w}$ | $\delta E_{D-I}$ | **-0.47** | **-0.45** |
| $\delta E_D$ | $\delta E_{D-I}$ | -0.02 | -0.01 |
| $\delta E_{D-w}$ | $\delta E_{D-I}$ | **-0.86** | **-0.82** |

This comparison speaks in favor of the rigidity of the ds-DNA and may explain the observed differences between a DNA and a protein system. In a similar fashion as before, we perform the Fourier transform of the energy autocorrelations and plot them in log-scale to find out the nature of 1/*f* noise character. The resultant plots are presented in Fig. 6.

**FIG. 6.**
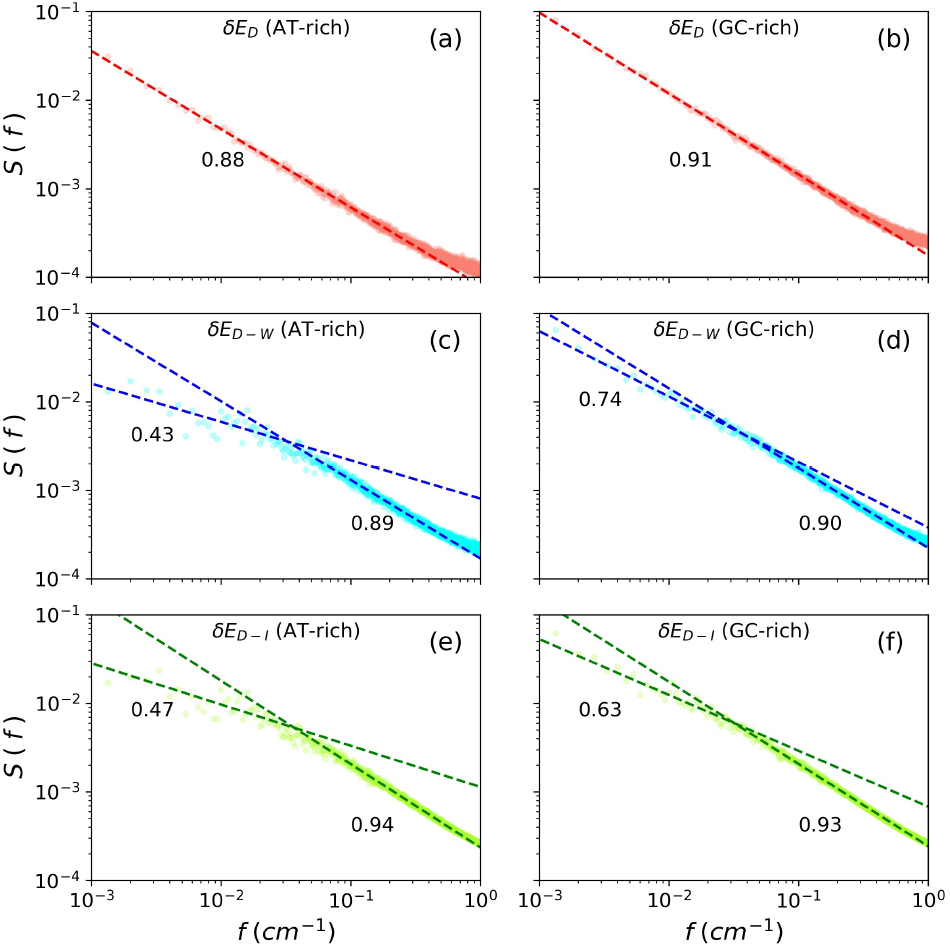
Spectra of energy fluctuations in ss-DNA solution (log-log representation). (a) DNA-self energy for AT-rich, (b) DNA-self energy for GC-rich, (c) DNA-Water cross interaction energy for AT-rich, (d) DNA-Water cross interaction energy for GC-rich, (e) DNA-Ion cross interaction energy for AT-rich ss-DNA, and (f) DNA-Ion cross interaction energy for GC-rich ss-DNA system. The numbers denote the slopes of the adjacent lines.

We observe that the unimodal 1/*f* noise character for DNA self-energy fluctuation and the bimodal character for DNA-Water and DNA-Ion cross interactions are retained in single-stranded DNA. However, we find an interesting correlation between DNA-water and DNA-ion interactions. Out of the two slopes in the 1/*f* noise spectrum, the slope with the lower of the two values increases and attain values larger than that of ds-DNA. On the other hand, the slope with the higher value (close to *∼*0.9) remains the same.

The smaller of the two exponents in this bimodal spectrum can be recognized as due to water. The interesting point in the increase in the value over that for ds-DNA is that the ss-DNA is more exposed to water and contains no stable grooves. Thus, the values tend to be closer to that of the exponent observed for neat water.

The larger value of the exponents that we attribute to the self-energy fluctuation of ss-DNA can arise from the intermittency of transitions in its energy landscape, or from the correlated dynamics with its surroundings. However, the exact nature of its origin is nontrivial to unveil and remains to be deeply analyzed.

### C. Dielectric relaxation and motion of counterions

The strong anti-correlation between in DNA-water and DNA-ion energy fluctuations suggests that the motion of the ions in the vicinity of DNA could also be driven by water. This in itself is not surprising but in the present case, ions are more localized (at least a fraction of them) around the negatively charged phosphate backbones that serve as a trap (energy minima) for the positively charged counter ions. So, a natural motion of the localized ions would transition from one trap to another trap. The strong anti-correlation observed points towards the possibility that such transitions could be mediated by water fluctuations. It is to be noted that such jump dynamics of the ions have directly been observed from an earlier MD simulation study.

Such systematic migration of counterions can give rise to a slow polarization relaxation that would respond to external electric fields. In fact, Oosawa proposed this motion of ions as a model of dielectric relaxation in aqueous DNA solutions.^17^ He envisaged a quasi-one-dimensional interacting random walk of ions along the DNA backbone. This is discussed in detail below. The anti-correlation observed here is related to this random walk and water fluctuations can provide the time constant involved in the said random walk.

We start with the exact expression for the frequency-dependent dielectric function of a dipolar liquid that is related to the total dipole moment time correlation function by Eq. 2.

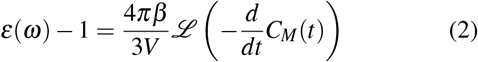

Here, *C*_*M*_(*t*) = ⟨**M**(0).**M**(*t*)⟩, *β* = 1*/k*_*B*_*T*, **M**(*t*) is the dipole moment vector of the system at time t, and ℒ denotes Fourier-Laplace transformation with respect to time with frequency *ω* as the conjugate variable. In the presence of free ions, the frequency-dependent dielectric function undergoes substantial modification,^29^ and the situation becomes more complex in the presence of a charged polymer with counterions, such as DNA. Here we detail the results (obtained by Oosawa) for the real (*ε*^′^(*ω*)) and imaginary (*ε*^″^(*ω*)) parts of the frequency-dependent dielectric constant [Eq. 3 and 4].

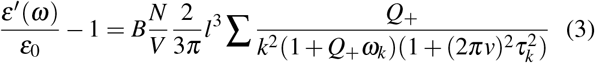

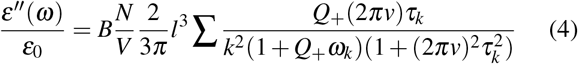

Oosawa derived these expressions for rod-like polyions. Here, *ε*_0_ is the static dielectric constant of the solvent, *N/V* is the number density of the polyion chain, *Q*_+_ is the density of bound counterions (approximated to be time-independent), *l* is the length of the polyion chain, *τ*_*k*_ is the relaxation timescale associated with the k^t^h mode, *ν* is the frequency, and 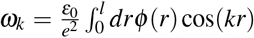, with *ϕ* (*r*) being the potential function. The model describes fluctuations in the position of the counterions along the chain in terms of polarization fluctuations that are Fourier decomposed. The time constant that enters is essentially a diffusion constant of counter ions along the chain.

The total frequency-dependent dielectric response is given by Eq. 5

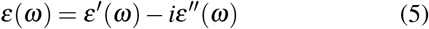

Although we start with the result of the model system of Oosawa to represent our aqueous DNA system, we introduce a waiting time distribution to generalize the random walk model. This in turn gives rise to a power law decay of polarization fluctuation which could be related to the 1/*f* noise discussed before.

The two phosphate backbones of a ds-DNA can be thought of as two polyion chains with almost equally spaced negatively charged traps where a fraction of the sites are blocked by the oppositely charged counterions. These trapped counterions possess a finite residence time around the DNA and move along the backbone from one site to the other, either influenced by the polarization fluctuation around DNA or as a random walker. This phenomenon leads to a fluctuation in the counterion population and generates a current around DNA, which contributes to dielectric relaxation. Restricted motion of counter ions along the DNA chain can give rise to a slow response.

The present approach is motivated by the study of Scher and Lax who explained the power law time dependence of current observed in plates after radiations.^30,31^ The main idea was that the electrons generated by optical flash execute an interacting random walk described by a waiting time distribution. In our model, the counterions can experience three types of transitions along the charged sites on the DNA chains, (i) counterions arriving from bulk to a binding site, (ii) ions leaving a binding site and diffusing to the bulk, and (iii) ions moving along the chain to another binding site, provided that is empty. These processes give rise to a distribution of waiting times, *ψ*(*t*).

Montroll et al. considered the flow of ions as the propagation of Gaussian current packets.^31^ The propagation velocity of the mean position of the packet, *l*, yields current. Thus, the generated electric current, *I*(*t*), becomes proportional to the time derivative of the solvation energy, *E*(*t*) [Eq. 6].

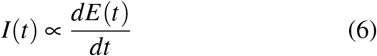

Now, movement from one site to another is associated with several factors such as potential barriers, availability of empty sites, and separation distance between sites. Moreover, there is spatial dispersion in the location of empty sites and potential barrier that depends on the local surroundings. Furthermore, there could exist bottlenecks on the path of migration of ions because the adjacent sites can be occupied, making an ion wait for a long time for the next jump.

Hence, one cannot approximate *ψ*(*t*) as a rapidly decaying function with a single transition rate. Following Scher and Lax, we approximate *ψ*(*t*) as 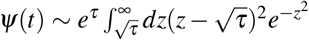 such that the long–time tail acquires a *τ*^−(1+*α*)^ dependence, where *α* lies between 0 and 1. We now define the probability of finding a random walker at position *l* at time t (*P*(*l, t*)) by the well-known Montroll-Weiss formula shown in Eq. 7.

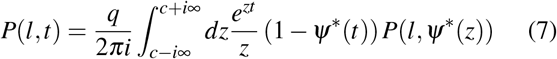

where, 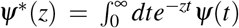 and *P*(*l, z*) is the random walt generation function defined as 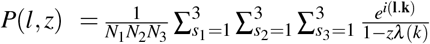, *k* = 2 *πs*/*N* and *λ* (*k*) is the hopping transition probability in the Fourier space, expressed as *λ* (*k*) = ∑_*l*_ *p*(*l*)*e*^*i*(**l.k**)^. Here, ∑_*l*_ *p*(*l*) = 1.

Given the probability and waiting time distribution the electric current exhibits power law as in Eq. 8.

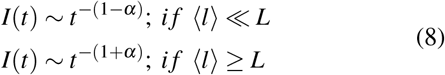

Here *L* is the total length of the DNA backbone. This agrees with the power law behavior (*∼t*^*α*^ or *∼t*^−*α*^) that has been observed in dielectric relaxation of DNA aqueous solution.

There however exist a relatively less number of detailed studies on DNA solutions compared to the same for protein solutions. The latter was pioneered by Pethig, Grant, and others. Here the situation is far more complex but probably a lot can be gained by carrying out such studies.

### D. DNA solvation dynamics

Power law decay is also exhibited by DNA solvation dynamics. Extensive work by Berg, Sen and co-workers have revealed that the time-dependent solvation of a dye attached to a DNA duplex showed a long-time power law decay such that the solvation time correlation function (TCF) survived beyond 100 ns.^14–16^ Such long-time decay was not expected a priori and a satisfactory explanation of this anomalous decay has defied several efforts. In the present study, we present two analyses. First, we use an already established relationship between the solvation time correlation function and dielectric relaxation function to explain the origin of the long-time decay. Second, we discuss the relevance of the observed DNAion-water dynamics discussed above in the measured solvation TCF.

In a continuum model description where dielectric relaxation is given by a single Debye relaxation time *τ*_*D*_, the solvation time correlation function is exponential, with the solvation relaxation time given by the well-known longitudinal time defined by *τ*_*L*_ = (*ε*_∞_*/ε*_0_) *τ*_*D*_^18^. This nice expression shows that solvation can proceed at a time scale much faster than the Debye dielectric relaxation time, *τ*_*D*_. However, this simple relation breaks down in systems where dielectric relaxation derives contributions from several components, like DNA, water, and ions. As already discussed in detail above, in aqueous DNA solution, dielectric relaxation display non-exponential power law dynamics which could originate from motion of counter ions along the DNA chain.

Nevertheless, a quantitative translation of dielectric relaxation data to predict solvation time correlation function is still possible, as discussed by Castner et al.^32^ Let us assume that the dielectric relaxation is non-Debye and given by the following expression [Eq. 9].^33,34^

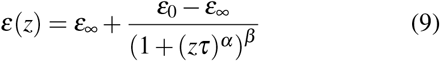

Eq. 9 is known as the Havriliak-Negami equation^35^ where *ε*(*z*) is the frequency-dependent dielectric function, *α* and *β* are parameters which lie between 0 and 1, *ε*_0_ and *ε*_∞_ are respectively the zero and infinite frequency dielectric permittivity of the solvent, and t is the relaxation timescale. Davidson-Cole formula (where *α* is set to 1) was previously used to study the dynamics of the reaction field^33^. For a dipolar solute, the time-dependent solvation energy (Δ*E*(*t*)) is given by Eq. 10

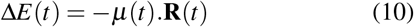

here *μ*(*t*) is the time-dependent dipole moment and **R**(*t*) is the reaction field. **R**(*t*) in the frequency domain can be expressed as *R*(*ω*) = *r*(*ω*)*μ*(*ω*) where the reaction field prefactor *r*(*ω*) is given by Eq. 11.

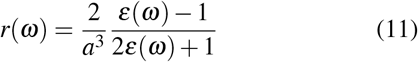

Here *a* is the cavity radius. It was shown that *r*(*t*) is a single exponential function when *β* = 1. For *β <* 1, *r*(*t*) becomes non-exponential. In such situations, a slow dielectric relaxation component gets translated to a slow solvation time scale. Because of the complex analytical forms involved, a closed-form solution of the solvation TCF is no longer possible, but a calculation can be carried out through Laplace transformation. To explain the occurrence of the power law decay, we note that in general the solvation time correlation function [*ϕ* (*t*)] can be expressed as the summation in Eq. 12.

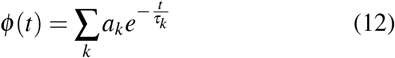

From the Oosawa model, we know that *τ*_*k*_ ∝ (1*/k*^2^). If we transform Eq. 12 into an integral and substitute the expression of *t*_*k*_, we obtain Eq. 13.

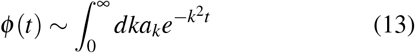

Eq. 13 in one- and two-dimensions gives *t*^−1*/*2^ and *t*^−3*/*2^ dependence of *ϕ* (*t*) respectively. Therefore, the origin of a power law decay can be explained within the Oosawa model. A detailed numerical solution is yet to be carried out.

Any quantitative study of solvation dynamics of a tagged dye (be it aminopurine or coumarin), must include all three contributing components: (i) DNA, (ii) water, and (iii) ions. Our present study showed that the last two components are themselves rather strongly anti-correlated (Table I). This gives rise to a scenario where a slow component can arise from the motion of the counter ions. As discussed above, solvation dynamics experiments might directly access the polarization relaxation due to the motion of the ions, thus explaining the slow component. Our studies reveal a slow component can arise from DNA energy fluctuations. We note that dielectric relaxation experiments probed long duplex chains while solvation dynamics experiments employ small chains. The length dependence of both solvation dynamics and dielectric relaxation phenomena remains unsolved.

### E. Role of a fluctuating environment in DNA functionality

One key aspect of DNA functionality, not discussed adequately in the literature, is the structural fluctuations essential to perform actions. This aspect has been discussed extensively for proteins.^7,8^ In Lehninger’s well-known textbook entitled “Principles of Biochemistry”, a section on DNA starts with the statement “DNA is a remarkably flexible molecule”.^36^ This flexibility is manifested in base pair opening, in the stretching of the length of a DNA segment, and in motions such as twist, rise, and roll of the adjacent base pairs. The detailed environment around DNA within the biological cell is not known precisely, in its restricted state wrapped around a histone octamer, although several recent studies have addressed this subject.^37,38^ In the present work we focus only on a free DNA molecule in an aqueous environment and attempt to learn about the fluctuations. It is conceivable that the fluctuations we discuss here partly survive within cells where an RNA polymerase allowed the formation of a transcription bubble. This is a complex process whose rate and mechanism have not been fully investigated.

We conjecture that a local, limited unwrapping of a ds-DNA can be induced if a certain configuration and/or arrangement of counter ions is formed by fluctuations. Such arrangements can exert force on the backbone. For example, the duplex can become locally unstable if on one side we have excess counterions while on the opposite side we have fewer numbers. This force imbalance can stretch or bend the duplex. In a real system within the cell, one would need to consider other factors like the role of enzymes that can facilitate bubble formation.^9,10^ But enzymes need to translate along the DNA.^39,40^ In such cases also, water and ion fluctuations must play an important role that has not been investigated. In an early enlightening study, Chandra showed that the presence of neighboring ions can accelerate the breaking of hydrogen bonds among water molecules.^41^ Such effects should be present not to any small degree in the present case and could play important role in base pair opening. Unfortunately, the present model and simulation timescale do not show base pair hydrogen bond breaking. We are working on this interesting problem.

## IV. CONCLUSION

The present study attempts to understand the coupling of single and double-stranded DNA with its immediate surroundings, by studying via simulations fluctuations in interaction energies between the DNA, water, and counterions. We believe that this is a novel approach as it can provide information about the energy landscape of the DNA-water-ion system.

Because of the complexity, the dynamics of this system have remained poorly understood. This is despite the existence of interesting experimental results in the areas of solvation dynamics and dielectric relaxation that display power-law decay in the long-time or low-frequency window. In both cases, the motion of counter ions along the negatively charged phosphate backbone of the ds-DNA is implicated, although no precise quantitative demonstration of the same has been put forward. The present study demonstrates that the motion of ions on the DNA backbone is coupled to water dynamics in a definite way. This cross-correlation is expected to contribute to the solvation dynamics of any probe dye placed in a DNA duplex.

Energy fluctuations have been previously shown to be efficient collective variables to understand the mechanism of the solvent control of protein dynamics.^7^ In the case of ds-DNA we find that the strength of coupling between fluctuations in DNA and water/ion is weak. However, a surprisingly large anticorrelation exists between the water and the counterions, manifested in the coupling of DNA-water and DNA-ion interaction energies. This indicates that the water molecules get displaced when a counterion approaches the biomolecule to bind with one of the available sites.

We hypothesize that the negatively charged phosphate groups present at regular intervals in the DNA strands serve as potential traps that regulate the movement of ions along the DNA backbone. A model based on continuous time random walk of these counterions along the DNA chains shows that the resultant current results in a power law of the waiting time distribution of the ions, which might explain the experimentally observed long-time tail in the solvation dynamics of DNA that can be fitted to a power law.

The observed low anticorrelation between ds-DNA self-energy and DNA-water interaction energy dramatically increases in the case of an ss-DNA. This could be a manifestation of the increased flexibility of the ss-DNA backbone, in absence of the base-pair hydrogen bonds that are present in a ds-DNA.

The power spectrum of the DNA energy fluctuations is characterized by 1/*f* noise properties. In the case of DNA-water and DNA-ion coupling energies, the power spectrum is found to be bimodal, where the two slopes resemble the self-energy fluctuations of DNA and water/ions. Surprisingly, the nature of the spectrum is indifferent to the flexibility of the DNA, such that we observe no significant differences between the spectra in the ds- and ss-DNA systems.

## ACKNOWLEDGMENTS

S.Mu. and S.Mo. thank the Indian Institute of Science (IISc) and Science and Engineering Research Board (SERB) for research associateship. BB thanks SERB (DST), India, for National Science Chair Professorship.

## Notes

### Competing Interest Statement

The authors have declared no competing interest.

